# Evaluation of SARS-CoV-2 isolation in cell culture from nasal/nasopharyngeal swabs or saliva specimens of patients with COVID-19

**DOI:** 10.1101/2023.02.16.528881

**Authors:** Shunsuke Yazawa, Emiko Yamazaki, Yumiko Saga, Masae Itamochi, Noriko Inasaki, Takahisa Shimada, Kazunori Oishi, Hideki Tani

## Abstract

It has been revealed that SARS-CoV-2 can be efficiently isolated from clinical specimens such as nasal/nasopharyngeal swabs or saliva in cultured cells. In this study, we examined the efficiency of viral isolation including SARS-CoV-2 mutant strains between nasal/nasopharyngeal swab or saliva specimens. Furthermore, we also examined the comparison of viral isolation rates by sample species using simulated specimens for COVID- 19. As a result, it was found that the isolation efficiency of SARS-CoV-2 in the saliva specimens was significantly lower than that in the nasal/nasopharyngeal swab specimens. In order to determine which component of saliva is responsible for the lower isolation rate of saliva specimens, we tested the abilities of lactoferrin, amylase, cathelicidin, and mucin, which are considered to be abundant in saliva, to inhibit the infection of SARS-CoV-2 pseudotyped viruses (SARS-CoV-2pv). Lactoferrin and amylase were found to inhibit SARS-CoV-2pv infection. In conclusion, even if the same number of viral genome copies was detected by the real-time RT-PCR test, infection of SARS-CoV-2 present in saliva is thought to be inhibited by inhibitory factors such as lactoferrin and amylase, compared to nasal/nasopharyngeal swab specimens.

## Introduction

The worldwide spread of the severe acute respiratory syndrome coronavirus 2 (SARS-CoV- 2) continues to mutate as the COVID-19 vaccine becomes more widespread. Currently, the real-time reverse transcription polymerase chain reaction (rRT-PCR) method, which can detect a specific gene of SARS-CoV-2, and the immunochromatography method, which can detect a specific antigen for SARS-CoV-2, are the two most popular methods mainly used for diagnosis of COVID-19^1^. Clinical specimens containing nasal swabs, nasopharyngeal swabs, sputum, saliva, etc. are used in these tests. It has been reported that the detection sensitivity in each specimen is highest for nasopharyngeal swabs, followed by nasal swab and saliva^2^.

We previously evaluated the viral isolation rate using cultured cells from clinical specimens positive for SARS-CoV-2, and analyzed the correlation with the Ct value of the rRT-PCR test^3^. However, although virus was efficiently isolated from the nasal/nasopharyngeal swab specimens, many copies could not be isolated from the saliva specimens. In this study, we compared virus isolation rates in clinical specimens of nasal/nasopharyngeal swabs or saliva that were brought to the Toyama Institute of Health for PCR testing for COVID-19. In addition, the viral isolation rates were evaluated for each variant of concern (VOC) mutant virus, including the omicron variants that are currently widespread worldwide. Furthermore, we also compared viral isolation rates by simulated specimens which contained cell culture-derived SARS-CoV-2 with the nasal/nasopharyngeal swabs or saliva samples of healthy donors who were not affected by COVID-19 and had not been vaccinated against the virus. Finally, we examined whether lactoferrin, amylase, cathelicidin, and mucin, which are known to be abundant in saliva, have inhibitory effects on SARS-CoV-2 infection. Elucidation of the infectivity of SARS-CoV-2 in saliva is expected to lead to measures to prevent infection in the future.

## Materials and Methods

### Cells and viruses

VeroE6 or VeroE6 cells overexpressing TMPRSS2 (VeroE6/TMPRSS2) (JCRB1819), which are considered to have high efficiencies of SARS-CoV-2 infection^4^, were obtained from the Japanese Collection of Research Bioresources (JCRB) Cell Bank (the National Institute of Biomedical Innovation, Health, and Nutrition, Osaka, Japan). The cells were grown in Dulbecco’s Modified Eagle Medium (DMEM; Nacalai Tesque, Inc., Kyoto, Japan) containing 10% heat-inactivated fetal bovine serum (FBS) and a 1% Penicillin-Streptomycin mixed solution. Cell culture derived SARS-CoV-2, Wuhan strain (hCoV-19/Japan/TY/WK- 521/2020, GSAID ID: EPI_ISL_408667), which was isolated at National Institute of Infectious Diseases (NIID) in Japan, was kindly provided from NIID.

### Specimens

Specimens of nasal/nasopharyngeal swabs or saliva suspended in phosphate-buffered saline or viral transport medium were collected from patients who were positive for SARS-CoV-2 by the rRT-PCR method performed at the Toyama Institute of Health from August 2020 to March 2022. Nasal/nasopharyngeal swabs (327 specimens) or saliva (268 specimens) were used for rRT-PCR and viral isolation tests, as anonymous samples. Differentiation of Wuhan strain or variants in each specimen was determined by rRT-PCR for the detection of the Alpha variant (N501Y), Delta variant (L452R), or Omicron variant (G339D) using a SARS-CoV-2 Direct Detection RT-qPCR Kit (TaKaRa Bio Inc., Shiga, Japan) with each specific primer/probe, or analysis by next generation sequencing. This study was approved by the Ethics Review Committee of the Toyama Institute of Health (R2-1).

### Viral isolation test from clinical specimens

Nasal/nasopharyngeal swabs or saliva specimens were stored at −80°C before being processed in cell culture. VeroE6/TMPRSS2 cells were inoculated with nasal/nasopharyngeal swabs or saliva specimens as described in an earlier report by Igarashi et al.^3^. Briefly, 20 μL of each nasal/nasopharyngeal swabs or saliva cleaning solution, which exhibited rRT-PCR positivity for SARS-CoV-2, was added to VeroE6/TMPRSS2 cells seeded the preceding day on a 24-well plate and cultured at 37°C for five days, and the cytopathic effect (CPE) was confirmed by visual observation under a microscope. Viral isolation was considered negative when the CPE was not observed for five days. All viral isolation procedures were performed in a biosafety level 3 laboratory at the Toyama Institute of Health.

### Viral isolation test from simulated specimens

The Wuhan strain of SARS-CoV-2 was amplified and its number of genomic copies and viral infectious titers were measured for undiluted lots. The viral solution was prepared to adjust to five different copies of viral genomes (5.0 × 10^2^, 5.0 × 10^1^, 5.0 × 10^0^, 5.0 × 10^-1^, 5.0 × 10^-2^ copies/μL) with nasal/nasopharyngeal swabs or saliva collected from 20 or 10 healthy donors who were not infected with SARS-CoV-2 and had not been vaccinated against COVID-19, as simulated specimens. Twenty μL of each simulated specimen was added to VeroE6/TMPRSS2 cells seeded the preceding day on a 24-well plate and cultured at 37°C for five days, after which the CPE was confirmed by visual observation under a microscope. Viral isolation was considered negative if the CPE was not observed after five days. All viral isolation procedures were performed in a biosafety level 3 laboratory at Toyama Institute of Health.

### Inhibitory test of saliva components on SARS-CoV-2pv infectivity

A luciferase assay using SARS-CoV-2 pseudotyped virus (SARS-CoV-2pv) was used to evaluate the infectivity of SARS-CoV-2. The Wuhan strain of SARS-CoV-2pv was generated using the vesicular stomatitis virus (VSV) pseudotyping system as described previously^5^. Lactoferrin, amylase, and mucin were purchased from Fujifilm Wako Chemicals, Osaka, Japan. Cathelicidin was purchased from Peptide Institute, Inc., Osaka, Japan.

To examine the effect of candidate substances on the cells, VeroE6 cells were transferred into 96-well plates the day before virus inoculation. The culture medium was removed, and 100 μL/well of each candidate substance adjusted for indicated concentration was added and incubated at 37°C for one hour. Next, 10 μL/well of SARS-CoV-2pv was inoculated and cultured at 37°C for one day. The value of relative light unit of luciferase was determined using the PicaGene Luminescence Kit (TOYO B-Net Co., Ltd., Tokyo, Japan) and GloMax Navigator System G2000 (Promega Corporation, Madison, WI, USA), according to the manufacturer’s protocol.

VeroE6 cells were treated with indicated concentrations of the substances for one day and cell viability was determined by measuring the luciferase activity using CellTiter-Glo 2.0 Cell Viability Assay (Promega) and GloMax Navigator System G2000, according to the manufacturer’s protocol. The percentage of reduction of viral infection and cell viability by the candidate substances were calculated using the luciferase activity, with the no-treatment control taken as 100%.

### Statistical analysis

Differences of viral isolation in clinical specimens and in simulated specimens were examined for statistical significance using a χ2 test (Fisher’s exact test was used when the expected frequency was less than five), and P<0.05 was considered significant. In the test of the inhibitory effects of saliva components, Dunnett’s test was used for comparison of all experiments, and P<0.05 was considered significant.

## Results

### Viral isolation from clinical specimens

For the study of SARS-CoV-2 isolation from clinical specimens in cell culture, 327 nasal/nasopharyngeal swabs and 268 saliva specimens were used. The CPE was confirmed in 149 of 327 nasal/nasopharyngeal swabs (45.6%) and 62 of 268 saliva (23.1%) specimens. It was found that the saliva specimens had a significantly lower isolation efficiency of SARS-CoV-2 than the nasal/nasopharyngeal swab specimens (Fig. 1A). In addition, when each specimen was compared by Ct value group, the viral isolation efficiency from saliva specimens was significantly lower in the group with Ct values of 20-25 [nasal / /nasopharyngeal swabs (NS): 81.3% vs saliva (S): 56.3%] and 25-30 (NS: 43.1% vs S: 18.8%), and no difference was observed in the group with Ct values of 30 or more (Fig. 1B). Subsequently, the viral isolation efficiencies were compared for each SARS-CoV-2 variant. The isolation efficiency from saliva specimens was significantly lower among the Wuhan strain (NS: 53.1% vs S: 21.8%) (Fig. 2A, left), Delta variant (NS: 46.7% vs S: 22.7%) (Fig. 2C, left), and Omicron variant (NS: 33.3% vs S: 8.5%) (Fig. 2D, left), but no difference was observed in the Alpha variant (NS: 52.0% vs S: 38.2%) (Fig. 2B, left). In addition, when each variant was compared by Ct value group, the isolation efficiencies of the Wuhan strain, Alpha variant, Delta variant, and Omicron variant were significantly lower than that of the nasal/nasopharyngeal swab specimens in the Ct value of the 25-30 group, 20-25 group, 25-30 group, and 20-25 group, respectively (Fig. 2, right panels). Furthermore, we compared viral isolation rates among the strains in the same specimen species (Fig. 3). In both nasal/nasopharyngeal swab and saliva specimens, the viral isolation rate was significantly lower for the Omicron variant than for the other variants. The Alpha variant showed a significantly higher isolation rate in saliva specimens compared to the other variants.

**Fig. 1.**
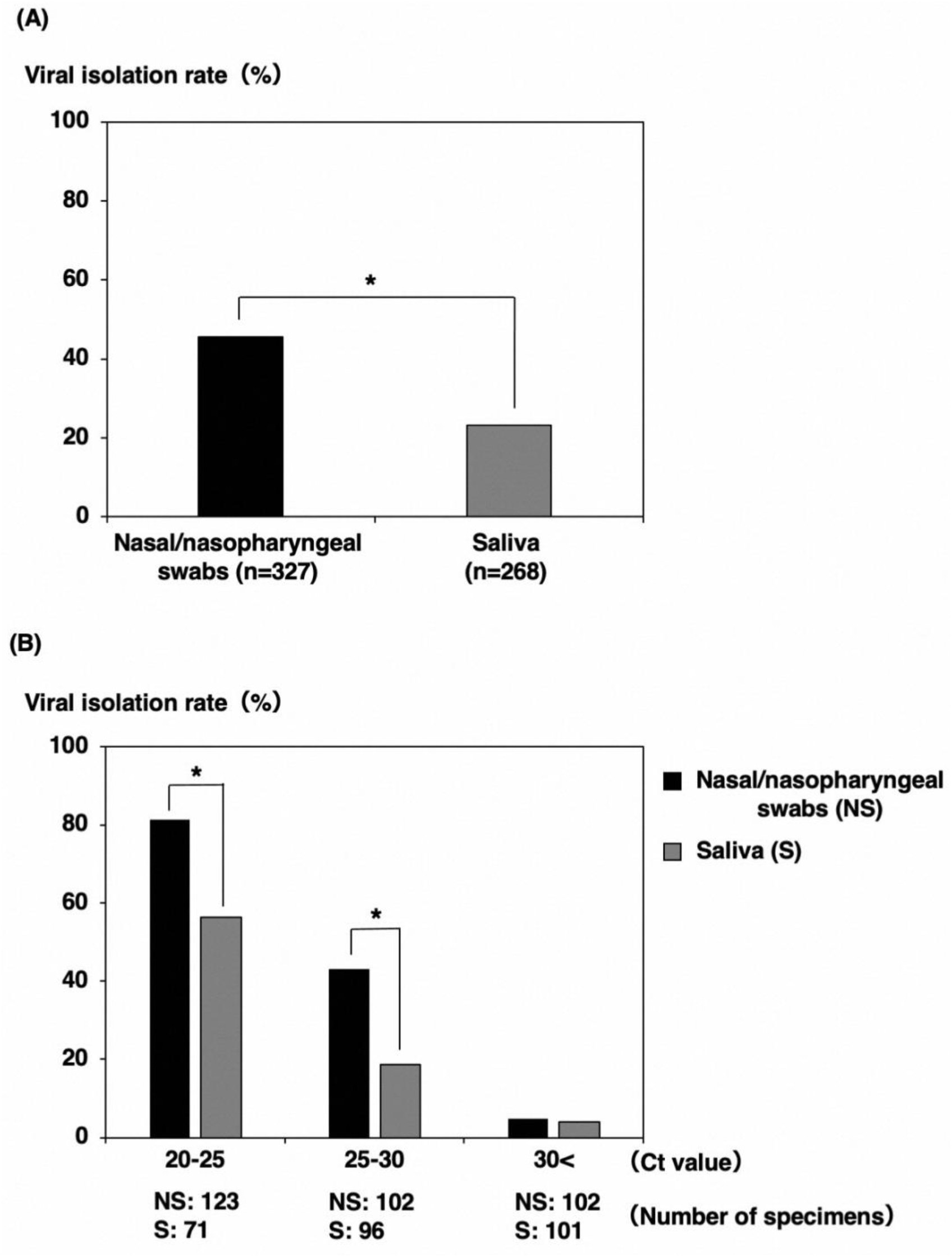
Comparison of viral isolation rate by clinical specimens. (A) Comparison of virus isolation rates between nasal/nasopharyngeal swabs and saliva. (B) Comparison of virus isolation rates by Ct value group (Ct 20-25, 25-30, 30<). *: *P*<0.05.

**Fig. 2.**
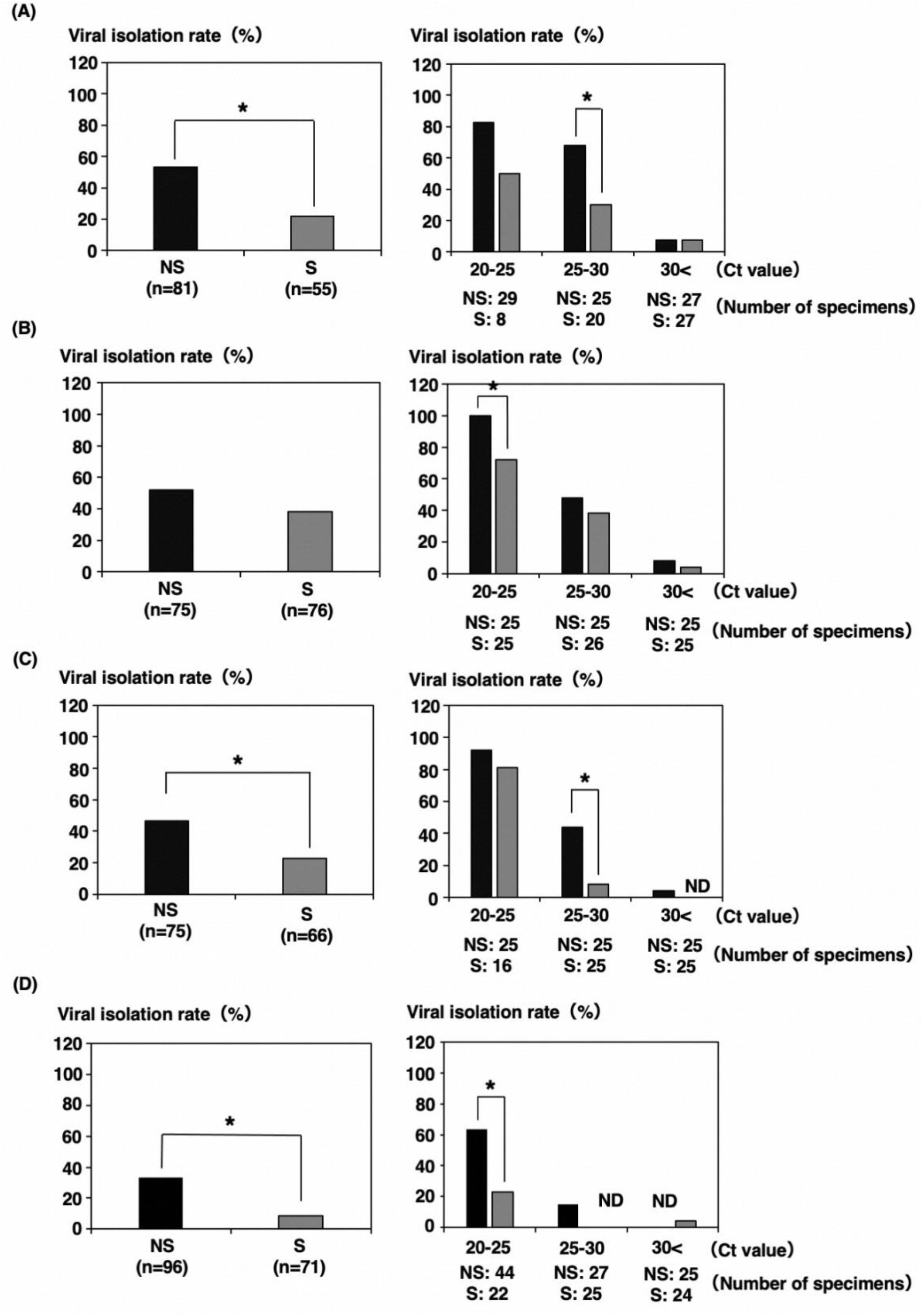
Comparison of viral isolation rates by SARS-CoV-2 variants [(A) Wuhan strain, (B) alpha variant, (C) delta variant, (D) omicron variant] between nasal/nasopharyngeal swabs and saliva. *: *P*<0.05.

**Fig. 3.**
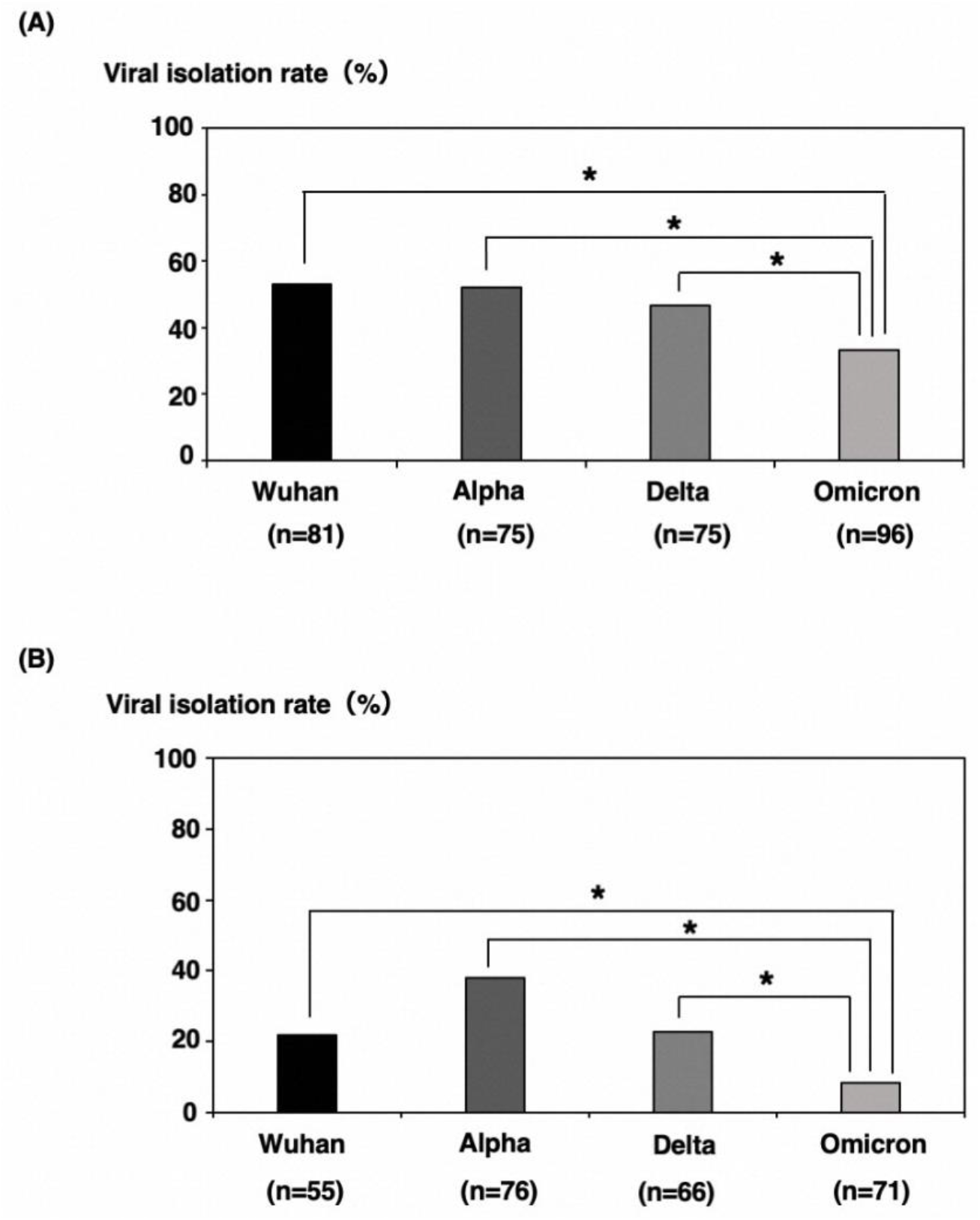
Comparison of viral isolation rates among virus strains in the same species of specimen. (A) nasal/nasopharyngeal swabs, (B) saliva. *: *P*<0.05.

### Viral isolation from simulated specimens

For the study of SARS-CoV-2 isolation using simulated specimens, the viral isolation efficiency from the saliva specimens was significantly lower than that of the nasal/nasopharyngeal swab specimens in 1.0×10^2^ copy [2.8×10^-1^ plaque forming unit (PFU)] (Fig. 4). In addition, there was no difference in over 1.0×10^3^ copy (2.8×10^0^ PFU) because all the nasal/nasopharyngeal swabs and saliva specimens could be isolated. On the other hand, both of the simulated specimens were not separated below 1.0×10^1^ copy (2.8×10^-2^ PFU).

### Effects of saliva components on SARS-CoV-2pv infectivity

Lactoferrin, amylase, cathelicidin and mucin, which are known to be abundant in saliva, were examined as for inhibitory components against viral infection in this study^6,7^. Lactoferrin treatment specifically inhibited SARS-CoV-2pv infection in a dose-dependent manner (Fig. 5A). Amylase treatment also significantly inhibited both SARS-CoV-2pv and VSVpv infection under a higher concentration (Fig. 5B). Meanwhile, cathelicidin treatment did not change up to 100 ng/mL, and was rather enhanced at 1,000 ng/mL in SARS-CoV-2pv infection (Fig. 5C). Mucin treatment showed no change in both SARS-CoV-2pv and VSVpv infection (Figure 5D). No cell cytotoxicity was observed in the treatment of each reagent within the range examined in this study.

**Fig. 4.**
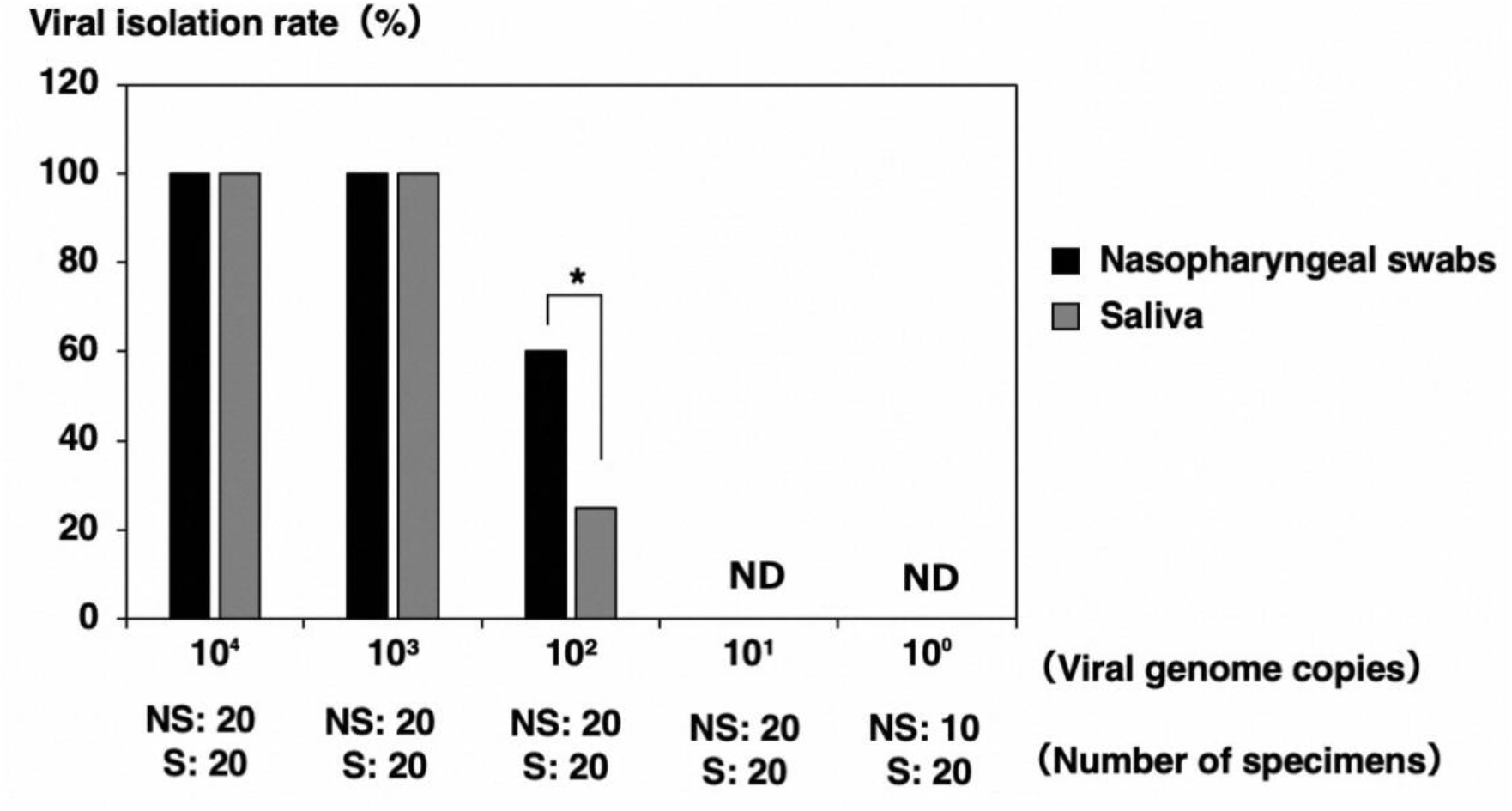
Comparison of viral isolation rates using simulated specimens. The virus used in this study was added by serially diluting the amount of the genome by 10-fold. ND: Not detected., *: *P*<0.05.

**Fig. 5.**
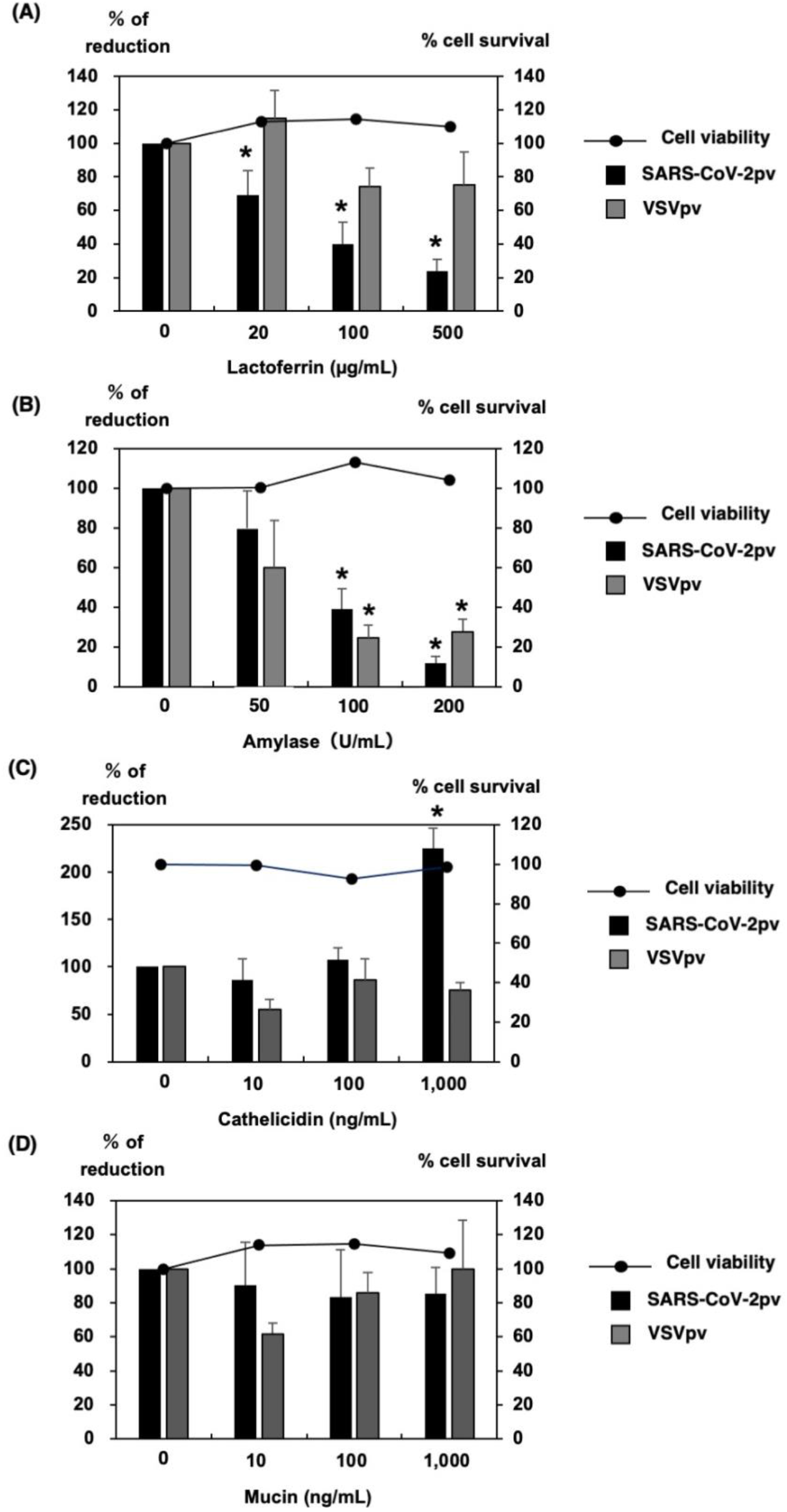
Effects of pseudotype viral infection by candidate substances. VeroE6 cells were infected with SARS-CoV-2pv or VSVpv and treated with indicated concentrations of lactoferrin (A), amylase (B), cathelicidin (C), and mucin (D). Data are presented as mean±standard deviation and repeated at least three times (n=3), *: *P*<0.05.

## Discussion

In this study, it was shown that the viral isolation efficiency in saliva specimens is significantly lower than that of nasal/nasopharyngeal swab specimens for COVID-19. As a result of analysis by Ct value, the viral isolation efficiency in saliva specimens was significantly lower in the groups with Ct values of 20-25 and 25-30, and no difference was observed in the group with Ct values of 30 or more. Therefore, it was found that the viral isolation efficiency differs depending on the type of specimens when the Ct value is 30 or less. In addition, previous studies have shown that the viral isolation efficiency decreases when the Ct value is 30 or more^3,8-11^. In the comparison of the viral isolation efficiency for each viral strain, the viral isolation efficiencies from the saliva specimens were significantly lower between the Wuhan strain, Delta and Omicron variants of SARS-CoV-2, but no difference was observed in the alpha variant.

In this study, specimens with low Ct values were not sufficiently collected among the saliva specimens containing the Wuhan strain and Delta variant of SARS-CoV-2. Therefore, it is possible that the viral isolation efficiency of the saliva specimens was significantly lower than the real isolation efficiency. On the other hand, the number of specimens could be the same in all Ct value groups, and there was no difference in the viral isolation efficiency between the nasal/nasopharyngeal swabs and saliva specimens for the Alpha variant of SARS-CoV-2. It will be necessary to increase the number of specimens to further verify whether there is a real difference among the variants. In the comparison of the viral isolation efficiency by Ct value group for each variant, the group with the significantly lower Ct value was different. However, although there was no significant difference, the viral isolation efficiency of saliva specimens tended to be low except for the groups with Ct values of 30 or more. Combined with the result of the comparison of the Ct value groups between variants, it is inferred that the viral infectivity decreases due to some interference by saliva components in the saliva specimens with Ct values of 30 or less. In a comparison of the viral isolation efficiency among the strain or variants in the same type of specimen, the viral isolation efficiency in the Omicron variant was significantly lower than that of the other strains or variants in both nasal/nasopharyngeal swab and saliva specimens. The Omicron variant has been reported to differ from other variants in terms of its dependence on TMPRSS2 during cell entry and the cleavage efficiency of viral spike protein^12^. Therefore, this may also affect viral isolation efficiency.

In the isolation of SARS-CoV-2 using simulated specimens, the viral isolation efficiency from the saliva-containing specimen was significantly lower than that of the nasal/nasopharyngeal swab-containing specimens in 1.0×10^2^ copy (2.8×10^-1^ PFU). The saliva used in the 1.0×10^2^ copy group was from patients 16-76 years old (median 42.5) and had a sex ratio of 7:3 (14 males and 6 females). The virus was not isolated in 15 of 20 saliva-containing specimens, and no significant difference was observed in age and sex compared to the five viral isolated specimens. All of the saliva used in this study was collected from healthy individuals who had negative SARS-CoV-2 PCR tests and had no history of vaccination against COVID-19, so the involvement of specific antibodies against the SARS-CoV-2 in saliva could be ruled out. Therefore, the decrease in infectivity of SARS-CoV-2 is considered to be due to the nature of saliva itself. By comparing the nasal/nasopharyngeal swabs and saliva samples using simulated specimens, the effect of viral interference by saliva on the clinical specimens in the previous experiment was experimentally shown.

Finally, we investigated which component of saliva was responsible for the lower isolation rate of saliva specimens by SARS-CoV-2pv. The results showed that among four candidate substances, lactoferrin and amylase inhibited SARS-CoV-2 infection the most. It has been reported that lactoferrin has an inhibitory effect on the entry and replication of SARS-CoV-2^13,14^. The concentration of lactoferrin in normal saliva is considered to be about 10 μg/mL^15^. Although that concentration is lower than what was used in this experiment, it is thought to have some anti-SARS-CoV-2 action in the saliva. In fact, it has been reported that human breast milk contains higher concentrations of lactoferrin than saliva, and that no infectious virus is present in breast milk even if the mother is infected with SARS-CoV-2^16^. Although the same experiment was performed with the control, VSVpv, for comparison, no significant inhibition was observed, although a concentration-dependent decreasing trend was observed. Therefore, it was suggested that lactoferrin has a higher inhibitory effect on SARS-CoV-2 infection. In addition, lactoferrin was not cytotoxic at the concentrations used in this study, suggesting that it may be useful as a preventive medicine against SARS-CoV-2 infection.

Amylase also inhibited SARS-CoV-2 infection in a dose-dependent manner at concentrations of 100 U/mL or higher. However, the concentration of amylase in normal saliva is reported to be about 30 U/mL^17^, which is much lower than the significant concentration used in this experiment. Therefore, it is thought that amylase has slight protective effect against *in vivo* infection. It also shows a similar inhibitory effect against VSVpv infection, and differs from the action of lactoferrin in that it is not specific for SARS-CoV-2. The reason for the decrease in the infectivities of both pseudotyped viruses is not well understood, but since amylase is an enzyme that degrades part of the sugar chain, it is thought that it has some effect on the viral spike protein or envelope protein for the entry. As for cathelicidin and mucin, neither SARS-CoV-2pv nor VSVpv showed an inhibitory effect for the infection. Rather, cathelicidin had an enhancing effect for the infection at high concentrations. From these experiments, it was considered that some substances contained in saliva had enhancing effects for SARS-CoV-2 infection, but that the inhibitory effect may have been largely derived from lactoferrin. In addition, although physiological concentrations of amylase alone cannot completely prevent SARS-CoV-2 infection, it is thought to synergize with lactoferrin to exert some anti-SARS-CoV-2 activity. It is assumed that these factors led to the lower isolation rates of saliva specimens compared to the nasal/nasopharyngeal swab specimens. Although this study examined four substances that are considered to be abundant in saliva, it is suspected that other substances that are specific to saliva may also be involved.

From these findings, it was speculated that infectious SARS-CoV-2 viral particles are more abundant in the nasal cavity and nasopharynx than in the saliva, even if the number of genome copies is the same as on the result of a rRT-PCR test. It was found that saliva acts in defensive manner with regard to infectious diseases, although not completely, and inhibits SARS-CoV-2 infection. In the future, it will be necessary to continue to study how these components of saliva act during SARS-CoV-2 infection.

## Author Contributions

Conceptualization and methodology, S.Y., K.O., and H.T.; sample collection, S.Y., E.Y., Y.S., M.I., N.I., and T.S.; investigation, S.Y., E.Y., and Y.S.; writing–original draft presentation, S.Y. and H.T.; writing–review and editing, S.Y., E.Y., Y.S., K.O., and H.T.; supervision, K.O. and H.T.; funding acquisition, H.T. All authors have read and agreed to the published version of the manuscript.

## Acknowledgments

We sincerely would like to thank Yoko Kanamori and Izumi Kawaguchi for technical and secretarial assistance.

